# DirectHRD enables sensitive scar-based classification of homologous recombination deficiency (HRD)

**DOI:** 10.1101/2024.06.21.600073

**Authors:** Ruolin Liu, Eugenia Roberts, Heather A. Parsons, Elizabeth H. Stover, Atish D. Choudhury, Justin Rhoades, Timothy Blewett, David D. Yang, Joyce F. Liu, Erica L. Mayer, Viktor A. Adalsteinsson

## Abstract

Homologous recombination deficiency (HRD) is a predictive biomarker for efficacy of PARP inhibition and platinum chemotherapy but remains challenging to detect from low tumor fraction samples such as liquid biopsies. Here, we describe DirectHRD, a whole-genome sequencing (WGS) scar-based classifier that is 10x more sensitive than state-of-the-art methods. DirectHRD can detect HRD at >=1% tumor fraction using 50x WGS of cell-free DNA.

## Main

Homologous recombination (HR) deficiency (HRD) is a clinically significant phenotype reflecting cells’ impaired ability to repair DNA double-stranded breaks (DSB). Cancer patients with HRD may benefit from synthetic lethal therapies such as PARP inhibitors (PAPRi) and platinum chemotherapies, but the detection of HRD remains challenging. Genomic scars caused by HRD have been used to identify patients who may benefit from PARPi^1,2^ and platinum chemotherapy. However, current HRD scar detection methods^3–7^ often require high tumor fraction (>20%) samples such as tissue biopsies; whereas, many cancer patients have much lower fractions of tumor DNA in plasma cell-free DNA (cfDNA).

Here, we present DirectHRD (Fig. 1a), an ultrasensitive scar-based classifier to detect HRD from low tumor fraction samples such as liquid biopsies using whole-genome sequencing (WGS) data. DirectHRD makes use of a specific type of genomic scar–a small, microhomology deletion (mhDel, Extended Data Fig. 1a)—believed to be the direct evidence of microhomology-mediated end joining in the absence of HR— due to its superior classification power^5,6^ and low error rate in next generation sequencing (NGS). However, mhDels represent only a fraction of already rare somatic small deletions. We thus surveyed PCAWG data^8^ for mhDels in 4 cancer types and found a median of 306 mhDels (151-537 for 15% and 85% quantile) in HRD-positive genomes defined by CHORD^5^ (Fig. 1b). We then modeled the number of mhDels that could be recovered at varied sequencing depths and tumor fractions and estimated that about 4-8 (15% quantile and median, respectively) mhDels could be uncovered at 1% tumor fraction using 50x WGS (Fig. 1c). This suggested that it could be possible, in principle, to utilize mhDels for HRD detection from low tumor fraction samples using moderate depths of WGS.

**Fig. 1:**
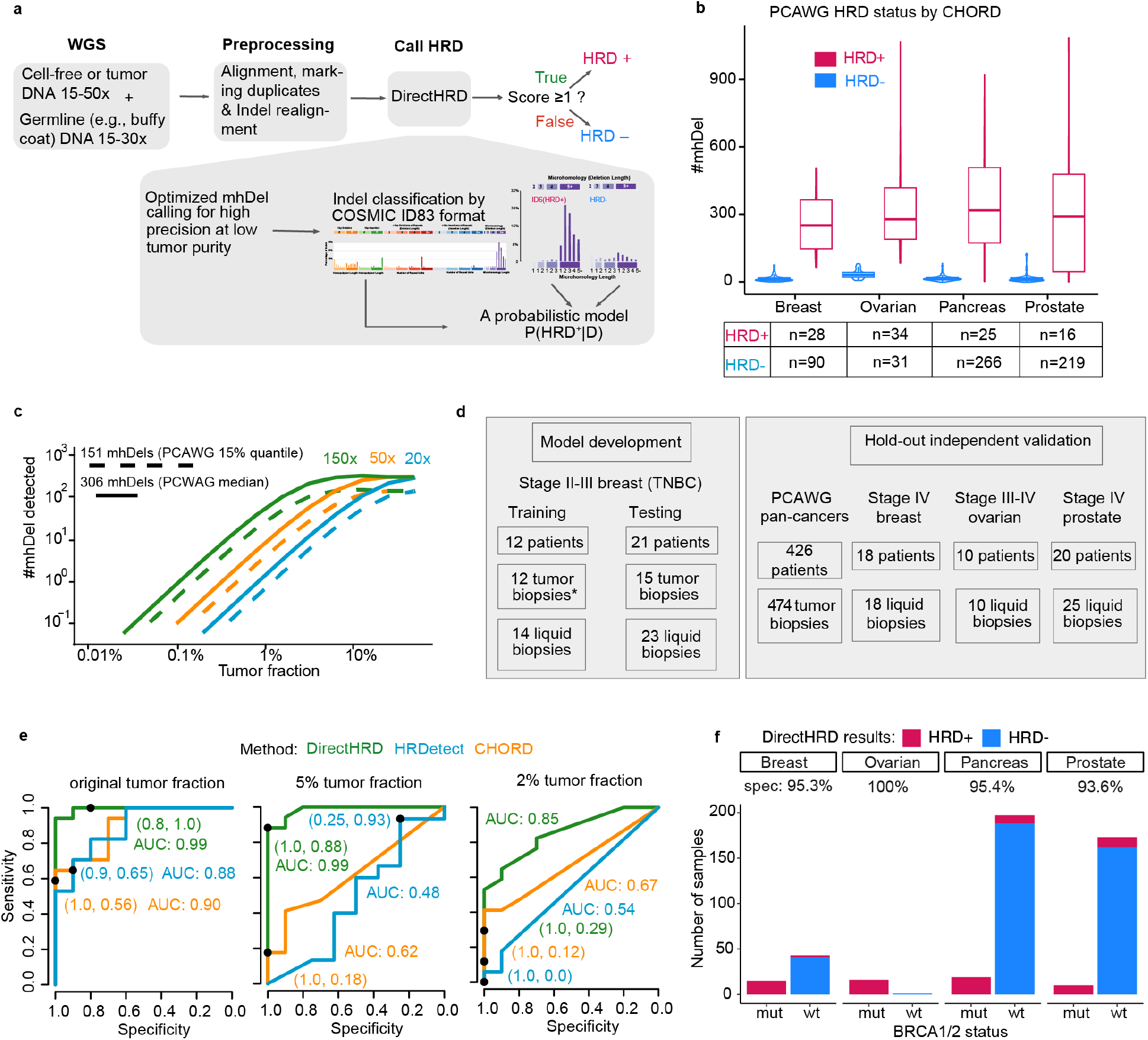
DirectHRD workflow and proof of concept. **a**, Schematic of DirectHRD. **b**, Overlay of boxplot and violin plots of number of microhomology deletions (mhDels) per sample in PCAWG data across 4 cancer types, stratified by HRD statuses calculated from CHORD. Center lines, boxes, and whiskers indicate medians, 25% and 75% percentiles, and 5% and 95% percentiles, respectively. The table below shows the number of HRD-positive and HRD-negative samples used in the plot. **c**, Modeling the expected number of mhDels with decreasing tumor fraction. Here, we use binomial models with 3 sequencing depths (20x, 50x, 150x) and 2 starting mhDel burdens corresponding to the 15% quantile (151) and median (306) number of mhDels in the PCAWG tumors classified as HRD-positive by CHORD. **d**, a summary of samples used in this study and their utilization for training, testing and hold-out validation. * indicates samples which were also sequenced with CODEC^9^. **e**, Receiver operating characteristic (ROC) curves and area under the curve (AUC) of HRD detection from tumor biopsies and simulated dilutions of tumor biopsies, left: undiluted 27 tumors downsampled at 15x coverage (tumor fractions: 14%-74%), middle: 5% tumor fraction at 15x, right: 2% tumor fraction at 15x. The black dot and the numbers in the parentheses indicate the performance (specificity, sensitivity) when the default cutoffs for HRD detection were used (DirectHRD score: 1.0, HRDetect p-value: 0.7, CHORD p-value: 0.5). **f**, DirectHRD results on 4 cancer types, including 474 PCAWG samples used by CHORD for performance evaluation, where the ground-truth HRD status was determined by bi-allelic loss-of-function status of BRCA1/2 genes, as detailed in Nguyen et al.^5^ (mut: mutant, wd: wildtype, spec: specificity).

We first developed DirectHRD using highly accurate CODEC^9^ technology, whole-genome sequencing of 14 tumor biopsies samples in TBCRC030, a previously published, prospective, randomized study of neoadjuvant paclitaxel vs. cisplatin in stage II-III triple negative breast cancer (TNBC)^10,11^. We found an HRD score cutoff of 1 in DirectHRD correctly classified tumors when using Myriad MyChoice HRD testing as ground truth (Extended Data Fig. 1b). However, we also recognized that standard Illumina WGS exhibits exceptionally low error rates for Indels (insertion/deletion polymorphism) in high complexity regions (versus other alterations such as single nucleotide variants, SNVs) and reasoned that it may be sufficient for mhDel detection (Extended Data Fig. 1c). Indeed, we adapted our CODEC single fragment caller (SFC) to standard Illumina WGS data and found that it detected mhDels with an error rate of 3×10^−9^ using a cutoff of 2 unique fragments (Extended Data Fig. 1c). Thereafter, we used standard Illumina WGS for DirectHRD. We showed superior performance for mhDel detection as compared to the state-of-the-art Indel detection methods (Extended Data Fig. 1d,e). After Indel calling, DirectHRD compares the Indel signature in each sample against a known HRD-positive signature (ID6 in COSMIC 3.2) and an HRD-negative signature which we derived from PCAWG HRD-negative breast cancers. We also derived an HRD-positive signature from PCAWG breast cancers to use as a backup signature (Methods). DirectHRD used this backup signature in 1/90 cfDNA sample and 1/81 synthetic or down-sampled tumor sample (Supplementary Table 6). Using an Indel signature such as ID6 allows us to assign posterior probabilities/confidence levels to each deletion and microhomology length combination (Extended Data Fig. 1f). We confirmed that the probabilistic model had better classification accuracy than using simple count feature (Extended Data Fig. 2a). Further details of DirectHRD are provided in Methods.

To validate DirectHRD on standard Illumina WGS, we first applied it to 15x WGS of tumor biopsies from 27 patients with stage II-III TNBC from the TBCRC030 trial^10,11^ including the 12 patients sequenced by CODEC (Supplementary Table 1), which we considered as the training data (n=12) and the remaining as test data (n=15, Fig. 1d). All 27 patients had tumor HRD status determined by Myriad MyChoice^3^ (Supplementary Table 1). When compared to this gold standard commercial assay^3^, we found DirectHRD outperformed CHORD and HRDetect^6^, two leading WGS-based classifiers^5^: AUC 0.99 vs 0.90 and 0.88 (Fig. 1e). Additionally, we compared DirectHRD to an orthogonal method, scarHRD^4^, which employs an HRD score^3^ from copy number and large structural variants similar to Myriad MyChoice. We found good correlation (0.81) between DirectHRD and scarHRD (Extended Data Fig. 2b). As an independent hold-out validation, we applied DirectHRD to 474 samples from the PCAWG project^8^, focusing on the four cancer types with the highest HRD prevalence (Fig. 1f, Supplementary Table 2). The ground-truth HRD statuses were determined by bi-allelic BRCA1/2 mutation assessment as established in Nguyen et al.^5^. We achieved 100% detection of patients with bi-allelic BRCA1/2 losses with high specificity (>90%) across all four cancer types.

Next, to explore the performance of DirectHRD at low tumor fractions, we created *in silico* dilutions of the same 27 tumor biopsies at 5% and 2% tumor fractions by diluting tumor WGS reads into matched normal WGS reads. DirectHRD maintained high performance at 5% (AUC: 0.99) and declined slightly at 2% (AUC: 0.85) (Fig. 1e). By comparison, CHORD yielded AUCs of 0.62 and 0.67 at 5% and 2% tumor fractions, and HRDetect resulted in AUCs of 0.48 and 0.54 at 5% and 2% tumor fractions, respectively. We noted that CHORD exhibited superior performance compared to HRDetect at lower tumor fractions. Consequently, we opted to exclusively benchmark against CHORD for cfDNA samples. Of note, we found CHORD’s performance improved using Strelka2^12^ instead of Strelka1^13^ as published for Indel calling. However, it still underperformed when compared to DirectHRD (Extended Data Fig. 2c). Finally, we confirmed that the performance held up in the test set after removing the 12 tumor biopsies that had also been previously sequenced by CODEC (Extended Data Fig. 2d).

To test the feasibility of detecting HRD from liquid biopsies, we sequenced 20 cfDNA samples from 10 HRD-positive patients and 10 HRD-negative patients with metastatic breast cancer^14^. The HRD statuses were found by the consensus results of scarHRD^4^ and SigMA^7^ on tumor whole-exome sequencing (WES) (Extended Data Fig. 3a-b). We matched selected cfDNA samples on cfDNA tumor fractions (Extended Data Fig. 3c) determined by ichorCNA^14^ and further refined the tumor fraction estimates using tumor WES-informed SNVs (Methods). We sequenced the 20 cfDNA libraries to a median 19x WGS coverage (range 4-27x, Supplementary Table 3). Two samples were excluded because the recalibration showed 0 tumor fraction (Extended Data Fig. 3d). DirectHRD correctly identified 9/10 HRD-positive cases (sensitivity 90%) and 7/8 HRD-negative cases (specificity 87.5%), while CHORD identified 2/10 HRD-positive samples (sensitivity 20%), both of which had tumor fractions >20% (Fig. 2). This is consistent with CHORD being suited for HRD detection in samples with higher tumor fractions. By comparison, DirectHRD missed 1 HRD+ sample with the lowest tumor fraction by WES-informed re-estimation (0.04%, Extended Data Fig. 3d). DirectHRD also falsely identified 1 HRD-negative sample as positive. Interestingly, this patient’s tumor biopsy had the highest scarHRD score at 39 among all HRD-negative patients, close to the cutoff 42 (Extended Data Fig. 3b). We further explored how the HRD scores compared between the cfDNA and tumor biopsy of each patient (Extended Data Fig. 3e) and found DirectHRD scores from cfDNA samples corrected for tumor fraction were correlated with scarHRD scores from the tissue biopsies (Spearman correlation *ρ*= 0.7).

**Fig. 2:**
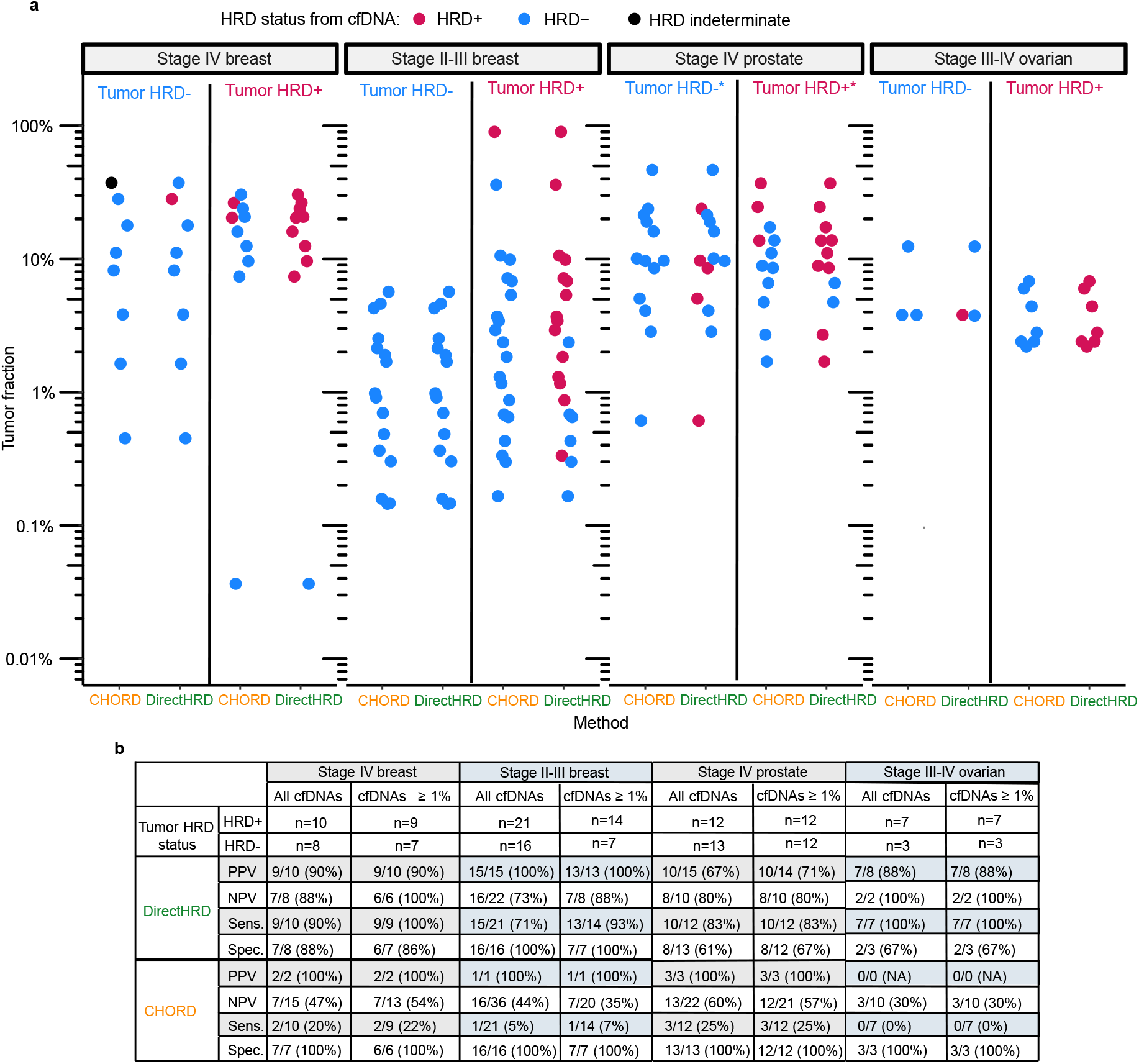
HRD detection from liquid biopsies. **a**, Comparison of DirectHRD and CHORD on (from left to right): 18 cfDNAs from 18 patients with stage IV (metastatic) breast cancer, 37 cfDNAs from 29 patients with stage II-III breast cancer (TNBC), 25 cfDNAs from 20 patients with stage IV (metastatic) prostate cancer, and 10 cfDNAs from 10 patients with stage III-IV ovarian cancer. Excepting prostate cancer, the HRD statuses inferred from patients’ tumors were used as ground truth and plotted side by side. For prostate cancer, the HRD statuses were inferred from germline or somatic pathogenic alterations in HR pathway genes from targeted panel sequencing of germline and cfDNA (indicated by *). HRD statuses from cfDNA by DirectHRD and CHORD were color-coded (magenta: positive, blue: negative). In stage IV (metastatic) breast cancer, CHORD classified 1 cfDNA sample as MSI-high and reported undetermined HRD status (black). **b**, Tabulated results of Positive Predictive Value (PPV), Negative Predictive Value (NPV), Sensitivity (Sens.), and Specificity (Spec.) of DirectHRD and CHORD.

Next, to explore the performance of DirectHRD on liquid biopsies from patients with stage II-III breast cancer, we utilized knowledge of the tumor fractions in cfDNA from prior testing using the MAESTRO MRD assay^11,15^, and selected 37 cfDNA samples from 29 patients with stage II-III TNBC from TBCRC030^10,11^ with tumor fractions of at least 0.1%. 92% of the cfDNA samples had tumor fractions <10%, and 43% had tumor fractions <1% (Extended Data Fig. 4a, Supplementary Table 1). 25 patients had HRD status from Myriad MyChoice, and we ran CHORD on the remaining 4 patients using their tumor WGS samples to determine their HRD statuses. We then performed WGS of all 37 cfDNA samples (median 43x, range 15-65.5x coverage) and applied DirectHRD and CHORD to all cfDNA samples. CHORD identified 1 cfDNA sample with 90% tumor fraction as HRD-positive and exhibited no false detection in the HRD-negative patients. DirectHRD correctly detected HRD down to ∼1% tumor fraction in cfDNA samples from HRD-positive patients, range (0.33%-90%), and with no false detection in cfDNA samples from HRD-negative patients (Fig. 2a). In all 37 cfDNA samples, DirectHRD achieved 100% (15/15) positive predictive value (PPV) and 71% (15/21) sensitivity. At ≥1% tumor DNA, DirectHRD yielded 93% (13/14) sensitivity (Fig. 2b). Similar performance was also seen within the 25 patients with MyChoice results for the first available cfDNA time point from each, mostly (n=23) at diagnosis (Supplementary Table 8). We additionally observed a high correlation (*ρ*=0.82, Extended Data Fig. 4b) between cfDNA and tissue HRD scores both from DirectHRD after correcting for tumor fraction. Since HRD score could be viewed as a quantitative measure of the HRD intensity, it suggests that DirectHRD on cfDNA could reveal the HRD intensity in a patient’s tumor biopsy. In assessing the impact of sequencing depth, we systematically downsampled the 37 cfDNAs to depths of 30x and 15x. Our analysis revealed comparable AUCs and specificities compared to the original data. However, there was a slight decrease in sensitivity at 30x (62% compared to 71% in the original data), with a further drop to 24% observed at 15x (Extended Data Fig. 4c). Lastly, we saw comparable results (Supplementary Table 9) after removing 14 cfDNA samples from 12 patients whose tumor biopsies had also been previously sequenced with CODEC and used in the original DirectHRD model derivation (the training set, Fig. 1d).

Lastly, we assessed DirectHRD’s performance on liquid biopsy samples in two additional cancer types. We sequenced 25 cfDNA samples (median coverage 47x, range 33-103x; median tumor fraction 9.7%, range 0.6%-46.6%) from 20 patients with metastatic castration-resistant prostate cancer (mCRPC) as previously published^16,17^(Supplementary Table 4) and 10 cfDNA samples (median coverage 58x, range 42-69x; median tumor fraction 3.8%, range 2.2%-12.4%) from 10 patients with stage III-IV ovarian cancer (Supplementary Table 5). Due to the lack of tumor tissue for mCRPC patients, we selected HRD-positive patients (n=10) if they had either germline or somatic pathogenic mutations in BRCA1/2 from targeted panel sequencing of cfDNA. We then identified HRD-negative patients if they did not have pathogenic mutations in BRCA1/2 and other HRR genes from targeted panel sequencing of cfDNA and selected 10 HRD-negative patients with similar cfDNA tumor fractions (Extended Data Fig. 5a,b). For ovarian cancers, we applied the same procedure to identify HRD-positive (n=7) vs HRD-negative (n=3) patients (Extended Data Fig. 5c,d) as we did in the metastatic breast cancer cohort. In comparing DirectHRD to CHORD for prostate and ovarian cancers, DirectHRD outperformed CHORD in sensitivity and negative predictive value (NPV) as shown in Fig. 2. For prostate cancer, DirectHRD achieved a sensitivity of 83% and NPV of 80% compared to CHORD’s 25% sensitivity and 60% NPV. Of note, neither DirectHRD nor CHORD detected HRD from the cfDNA of a BRCA1-mutant prostate cancer patient, and it is known that biallelic loss-of-function mutations are less common in BRCA1-compared with BRCA2-associated prostate cancers^18^, and thereby, this may not be a false negative. For ovarian cancer, DirectHRD demonstrated a sensitivity of 100% and an NPV of 100%, while CHORD’s sensitivity was 0% and NPV was 30%. DirectHRD’s specificities were lower in these two cohorts, possibly due to 1. the inability to use an established HRD scar classifier to establish ground-truth HRD status in the prostate cancer cohort for lack of tumor tissue (a previous study demonstrated that a substantial fraction of mCRPC with genomic features of HRD lacked biallelic loss of a core HRR-associated gene^19^), and 2. the small number of HRD-negative cases (n=3) in the ovarian cancer cohort. CHORD, on the other hand, had poor sensitivity when tumor fractions were low—25% when tumor fraction was between 10-20%, and 0% when tumor fraction was below 10% across both cohorts. Lastly, we found that the DirectHRD scores from cfDNA again correlated with the HRD intensities in patients’ tumor biopsies (Extended Data Fig. 5e).

In all, we showed that DirectHRD can detect HRD from low tumor fraction samples such as liquid biopsies with 10x greater sensitivity than the leading methods. We achieved this by identifying the best feature for HRD detection, mhDels, a well-defined scar of HRD. Compared to other scar-based methods, DirectHRD solely depends on scarce mhDels, and so we built on our experience developing CODEC to create an optimized deletion caller. Indeed, the novel aspects of this work include: 1) demonstrating that we can reliably detect mhDels in low-purity samples, such as liquid biopsies, using moderate depths of WGS with a custom caller; 2) proving that using mhDels alone is sufficient for HRD detection; and 3) developing a probability model where the Indel signature of HRD (Cosmic ID6), characterized by microhomology deletions, further improved our ability to detect HRD. We showed superior performance compared to the state-of-the-art methods which use all scar features. Our approach could enable detection of HRD in multiple cancer types and in early and late stages of cancer, including acquisition of HRD^20^, when tissue biopsies are insufficient or infeasible^10,21^. In the future, we plan to incorporate tumor fraction estimation into DirectHRD so that the patients with negative results and tumor fraction below our limit of detection could be recommended for other follow up testing. We also expect that higher sensitivity could be achieved with deeper WGS – the cost of which is plummeting – and that targeted sequencing for HR pathway mutations could be performed from the same cfDNA libraries as DirectHRD. Although DirectHRD showed high concordance with tissue-based HRD classifiers, additional studies will be needed to further establish the clinical validity and utility for HRD scar testing in multiple cancer types.

## Methods

### Indel classification and modeling number of mhDels in liquid biopsies

We downloaded the PCAWG Indel VCFs (Variant Calling Format) of 4 cancer types, Breast, Ovary, Pancreas, and Prostate from ICGC (https://dcc.icgc.org/). The HRD status of each sample was provided by Nguyen et al.^5^ in the supplementary table. We first restricted Indels to high complexity and high mapping quality regions (GIAB V3.0 easy regions, total 2.3B, https://ftp-trace.ncbi.nlm.nih.gov/giab/ftp/release/genome-stratifications/v3.0/GRCh37/LowComplexity/GRCh37_notinAllTandemRepeatsandHomopolymers_slop5.bed.gz) and then used the SigProfilerExtractor^22^ to categorize each Indel into one of the 83 classes according to COSMIC 3.2 (https://cancer.sanger.ac.uk/signatures/). There are 11 classes belonging to mhDels, and we summed all of them to get the total mhDel burden per sample. We took the median and 15% quantile of total mhDel counts in all HRD-positive samples across the 4 cancer types. We then used a binomial model where the success probability equaled half of the tumor fractions (assuming the mhDel is heterozygous) and the number of trials equaled the coverages. The probability of recovering an mhDel was calculated based on at least two successes.

### Indel calling

DirectHRD’s Indel caller is based on the single-fragment caller (SFC) from our previous work^9^. The SFC was described in detail in Bae et al.^9^. Some filters were implemented specifically for CODEC and were modified accordingly for standard NGS. Most importantly, we required a mutation to be supported by at least two unique fragments for error correction in the standard NGS data. The full list of parameters and command lines are given at https://github.com/broadinstitute/DirectHRD. Additional SFC Indel-filters (not in Bae et al.) include the following:

1. An Indel of interest is at least 8bp from the ends of a read.
2. No germline SNP or Indel is adjacent (within 5bp) to a Indel of interest.
3. Unique read depth of a matched germline at an Indel of interest location is >= 10
4. The read which contains an Indel of interest does not have another SNV or Indel nearby (for SNV ± 3bp, for Indel ± 10bp).

### Benchmarking Indel calling

We benchmarked the SFC Indel caller used in DirectHRD against established callers such as Mutect2^23^ and Strelka1^13^ and 2^12^ using a set of 6 positive control samples from the TNBC cohort^11^ and 4 negative control samples from healthy donors’ buffy coat DNA. For a fair comparison, we applied a high-complexity region filter to all callers, requiring that an Indel must occur in the high-complexity regions of the reference genome hg19 (GIAB V3.0 easy regions, total 2.3B). We included Strelka1 because it is the default Indel caller by CHORD, but we also tested the next generation caller (Strelka2) for comparison. For the positive control samples, we first selected 6 tumor WGS with the highest matched germline coverages and then downsampled the 60x tumor biopsies to 10x to mimic situations of low purities and looked for mutations that were called in both 10x and 60x. In the negative control data, we utilized technical replicates of germline DNA from 4 healthy donors. Each healthy donor had 2 WGS replicates sequenced in separate batches. The first batch had an average coverage of 16x (range 15-18x) and the second had an average coverage of 27x (range 23-30x). We used the higher (case) vs. lower (control) coverage data for each patient for paired “normal-normal” analysis.

### DirectHRD model and HRD score

The name ‘DirectHRD’ is inspired by a widely referenced review paper by McVey and Lee^24^ where they called microhomology-mediated end-repair (MMER) of a double strand break (DSB) as “director’s cut”. MMER as an alternative pathway to repair DSB creates unique mutation signatures, i.e., small deletions with microhomology, referred to as mhDel in this study (Extended Data Fig. 1a). Two COSMIC Indel signatures, i.e., ID6 and ID8, are found to be caused by repairing DNA DSBs by non-homologous DNA end-joining mechanisms. However, ID8’s etiology is less clear (as per COSMIC website, https://cancer.sanger.ac.uk/signatures/id/id8/) than ID6, and it has been found in a much higher prevalence and in tumor types that are less common to have HRD. Therefore, we chose to use ID6 as the HRD-positive signature and ID8 as a decoy signature. We also created an HRD-negative Indel signature using a public WGS dataset, PCAWG^8^: we first identified 66 breast cancer samples of 0 HRD probability according to CHORD’s prediction (CHORD Supplementary Data 1); we then downloaded the Indel VCFs from the ICGC portal (https://dcc.icgc.org) and summarized the Indel frequency according to the ID83 format from COSMIC. Similarly, we also created PCAWG HRD-positive signature from 27 breast cancer WGS samples of at least 75% HRD probability by CHORD. As expected, the cosine similarity between PCAWG HRD-positive signature and ID6 was high (93%) whereas it dropped to 73% when compared to ID8. The PCAWG HRD-positive signature will be substituted in if ID6 fails to converge in the EM algorithm. In ID83 format, an mhDel is subclassified by the deletion length and the length of the microhomology. There are 11 types of mhDel and the count vector of each mhDel type plus the deletion of 5 or more base pairs that is not repeat or microhomology mediated (5:Del:R:0) is used as the input of DirectHRD, which probabilistically assigns each mhDel to these signatures (e.g., ID6, ID8 or PCAWG Breast HRD-negative signature) using a multinomial mixture model (MMM). The score of each mhDel is essentially a posterior probability (thus 0-1) of being relating to HRD. The final HRD score of DirectHRD is the sum of scores of mhDels of ≥ 5bp deletion and ≥ 1bp homology. To be more specific, the likelihood of the MMM model is as follows:

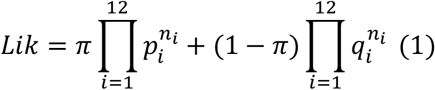

Here, *n*_*i*_ is the observed count of Indel feature type *i* and *π* is the mixing proportion and can be considered as the fraction of HRD-positive signature. *P, Q* are two multinomial distributions with parameters *p*_*i*_, *q*_*i*_ representing the probability of observing each count type from the positive and negative signatures, respectively. To maximize the likelihood function (1) with respect to *π, p*_*i*_, *q*_*i*_, we used Expectation-Maximization (EM) algorithm^25^. The initial values of *π* = 0.5 and *P, Q* are, respectively, probability vectors of ID6 and PCAWG Breast HRD-negative signatures. The EM algorithm iterates between taking the expectation of the log-likelihood evaluated at the current estimates of the parameters and a maximization step which calculates a new set of parameters. Since we have more parameters than data, and to avoid overfitting the data, we added a decay parameter *α* = 1/*N*_*mhDel*_ when updating *P, Q* : for example, *P*^*j+*1^ = (1 − *α*)*P*^*j−1*^ + (*α*)*P*^*j*^, and similarly, for Q. Here, *j* indexes the iterations. Intuitively, *P, Q* should be close to the signatures derived from a large population, and this decay parameter *α* slows down the change of *P, Q* from their initial values. The iteration is terminated if the change of *π* between two consecutive iterations is less than 10^−5^. In practice, we often observed the number of iterations is < 30. We set the default maximum number of iterations is 100. After obtaining *P, Q and π*, we calculated the posterior probabilities of mhDel type *i* by: *p*_*i*_*π*/(*p*_*i*_*π* + *q*_*i*_(1 − *π*)). In the case of a third signature being used, i.e. ID8, we introduced a third set of parameters *J or j*_*i*_ and the parameter *π* became a vector of *π*_*p*_, *π*_*q*_, *π*_*j*_ with a constrain *π*_*p*_ +*π*_*q*_ + *π*_*j*_ = 1.

### *In-silico* down-sampling and simulated dilution of tumor biopsies

In order to understand DirectHRD performance on lower tumor content samples, we first constructed *in silico* dilution series of tumor WGS from the stage II-III breast cancer (TNBC) cohort^10,11^. Due to the limit of sequencing depths of matched germline controls (∼15x, Supplementary Table 1) (which effectively limited the depths when creating low purity *in silico* tumors), we chose to have uniform coverage of ∼15x for all *in silico* tumors at various tumor fractions (undiluted, 5% and 2%). To do so, we downsampled the tumor and normal from each patient separately based on the targeted depth and tumor fraction. For tumors, the downsample fraction was calculated as (15x * targeted_tumor_fraction) / (original_tumor_depth * original_tumor_fraction). For normals, the downsample fraction was calculated as (15 – tumor_downsample_fraction * original_tumor_depth) / (original_normal_depth). Here, the original_tumor_fractions were calculated by Absolute^26^. To be consistent, the 15x downsampled undiluted samples instead of the original samples were used consistently throughout this study for calculating HRD scores from tumor biopsies in the TNBC cohort.

### Preprocessing WGS samples for HRDetect and CHORD

Both HRDetect and CHORD require pre-computed variant files including SNVs, Indels, and SVs (HRDetect also requires allele-specific copy numbers). Both papers published their own preprocessing workflows for obtaining these variant files. We followed exactly Nguyen et al. and Davies et al. for their variant calling workflows. In a nutshell, Strelka1 and Gridss1 were used in Nguyen et al. for small and large variants respectively. The Sanger institute’s WGS pipeline (available as a Docker file at https://github.com/cancerit/dockstore-cgpwgs) was used in Davies et al.

### Sample selection and establishing patients’ HRD statuses in the three independent cfDNA validation cohorts

In the metastatic breast cancer cohort^14^, we established the ground-truth HRD status from patients’ tumor WES. We ran two orthogonal methods and took the consensus results of the two methods. scarHRD^4^ sums a HRD score from large-scale variations including Loss of Heterozygosity (LoH), Large State Transition (LST) and Telomeric Allelic Imbalance (TSI). Whereas, SigMA^7^ uses Signature-3 from COSMIC, i.e., the HRD signature, which is derived from point mutations. To run SigMA, we first ran MutectClick or tap here to enter text.^27^ on the tumor-normal WES pairs to get the list of somatic SNVs, which SigMA used as input to its multivariate analysis (MVA) model. Two levels of confidence are output by MVA: MVA-pass and MVA-pass-strict. The latter represents a more stringent cutoff for HRD positivity. We first found 14 out of the 99 patients have at least 1 tumor sample that was HRD-positive by SigMA (MVA-pass-strict). However, 3 out of the 14 had inconsistent results from different tumor samples from the same patient. The remaining 11 patients were further tested by scarHRD. One patient did not meet the cutoff of HRD positivity (42) and was thus excluded. Since there were a lot of HRD-negative patients by SigMA (n=85) (expected from an ER*j* breast cancer cohort), we randomly selected 35 out of them and ran scarHRD on their tumor samples. 9 patients had at least one HRD-positive tumor sample according to scarHRD (cutoff 42) and were thus excluded. At last, we manually selected 10 out of the remaining 26 negative patients whose tumor fractions of the first available liquid biopsies matched those from the 10 selected HRD-positive patients (REMARK diagram in Extended Data Fig. 3a). The tumor fractions of the cfDNAs were estimated from ultra-low-pass whole-genome sequencing (ULP-WGS) using ichorCNA^14^. All WES and ULP-WGS samples were sequenced as part of a previous study^14^.

The prostate cancer patients included in this study were previously analyzed for associations between HRD status and clinical benefit from radium-223^16^ and copy number changes detected by ULP-WGS and response or resistance to docetaxel^17^. All patients had metastatic castration-resistant prostate cancer (mCRPC), primarily to bone, so tumor biopsies were not routinely performed. Instead, ULP-WGS of cfDNA was carried out and targeted sequencing of cfDNA was performed using an institutional prostate cancer-specific panel of 319 genes with duplex sequencing^28^. Germline and somatic variants were called using DeepVariant^29^ and Mutect2^23^, and these variants were annotated with Funcotator^30^. We identified 6 out of 138 patients in the radium-223 study and 4 out of 81 patients in docetaxel study with known BRCA1/2 pathogenic or likely pathogenic mutations (by ClinVar categories 4/5, Supplementary Table 4), categorizing them as HRD-positive patients. Patients with any pathogenic or likely pathogenic mutations in other HRR genes (RAD51, PALB2, ATM, CHEK2, CDK12, BRIP1) were excluded, resulting in the exclusion of 15 patients from the radium-223 study and 1 patient from the docetaxel study. From the remaining patients, we randomly selected 6 patients from the radium-223 study and 4 patients from the docetaxel study as HRD-negative patients (Extended Data Fig. 5a). We visually and statistically compared the tumor fractions of cfDNAs from all HRD-positive and HRD-negative patients. This random selection process was repeated until there was no significant difference between the two groups (Extended Data Fig. 5b).

In the ovarian cancer cohort, we first identified 39 patients whose tumor and plasma specimens were collected for another study (Han and Stover et al., under review). We then excluded patients if their blood normal, tumor, or cfDNA samples pre-treatment had not been sequenced. This filtering left us with 11 patients. Similar to the metastatic breast cancer cohort, we determined the ground-truth HRD status from patients’ tumor WES using the consensus results of scarHRD and SigMA. One patient was further excluded due to inconsistent HRD results between the two methods, resulting in a final cohort of 10 patients, of whom 7 are HRD-positive (Extended Data Fig. 5c,d).

### Estimation of cell-free DNA tumor fractions in three independent validation cohorts

In the metastatic breast cancer cohort, we selected samples with both tumor WES and cfDNA ULP-WGS^14^. We first used the cfDNA ULP-WGS tumor fractions estimated by ichorCNA to select cfDNA libraries for deeper WGS. Due to ichorCNA’s limit of detection of 3% tumor fraction on ULP-WGS, we recalibrated the tumor fractions by tracking tumor-informed SNVs in the deeper WGS cfDNA libraries. To do this, we first derived a list of somatic SNVs (we called fingerprints) from each patient’s tumor-normal WES data, a.k.a. the Mutect call from the previous section. We then used MIREDAS^31^ to filter reads and calculate variant allele frequencies (VAFs) on the fingerprints and multiplied the mean VAF by 2 to derive the recalibrated tumor fractions.

In both prostate cancer and ovarian cancer cohorts, the target panel sequencing of cfDNAs was performed. Tumor fractions were estimated using the mean variant allele frequency of somatic mutations in each sample.

### Comparing HRD scores between tumor and liquid biopsies

When comparing the HRD scores from cfDNA samples and tumor biopsies of the same patients, we used the raw HRD scores in the tumor biopsies and adjusted the scores by the tumor fractions in the cfDNA samples. The adjusted scores represented the expected HRD intensities from cfDNA and were calculated by simply multiplying the raw scores from the tumor biopsies by the cfDNA tumor fractions in the same sample. In the metastatic breast cancer and ovarian cancer cohorts, the raw tumor HRD scores were calculated from WES by scarHRD. In the TNBC cohort, the raw tumor HRD scores were calculated from WGS by DirectHRD and scarHRD.

### Statistics and reproducibility

The sample size was determined by the availability of tissue biopsies in each cohort, the number of HRD-positive patients, and the cfDNA tumor fractions. No statistical method was used to predetermine the sample size. The experiments were not randomized. The investigators were not blinded to allocation during experiments and outcome assessment. All statistical analyses in this work can be reproduced by codes on our GitHub repository https://github.com/broadinstitute/DirectHRD.

## Data availability

DNA sequencing data generated for this will be deposited into the controlled access database, such as dbGaP. GIAB easy region V3.0 is available at https://ftp-trace.ncbi.nlm.nih.gov/giab/ftp/release/genome-stratifications/v3.0/GRCh37/LowComplexity/GRCh37_notinAllTandemRepeatsandHomopolymers_slop5.bed.gz

## Code availability

The SFC/mhDel caller is available at https://doi.org/10.5281/zenodo.7630357 and https://github.com/broadinstitute/CODECsuite. DirectHRD and code required to reproduce the analyses is available at https://github.com/broadinstitute/DirectHRD.

## Acknowledgements

The authors acknowledge the Gerstner Family Foundation for its generous support. We are grateful for the funding support to the TBCRC from The Breast Cancer Research Foundation and Susan G. Komen. We thank Laurel Walsh and Rachel Li for their help with data visualization. We also thank Ofir Cohen for providing data access to the metastatic breast cancer cohort. Finally, we thank Zoltan Szallasi and Mary Lynne Hedley for their helpful comments on the manuscript.

## Author contributions

V.A.A and R.L. conceived and designed the project. R.L. and V.A.A developed the method. H.A.P., A.D.C, E.H.S, J.F.L, and E.L.M collected DNA samples. D.D.Y. performed analysis of targeted sequencing for prostate cancer samples. E.R. performed experiments. R.L. analyzed the data. R.L., V.A.A., E.R., H.A.P., A.D.C., E.H.S., E.L.M., J.R., T.B. interpreted the data. R.L., V.A.A., H.A.P., A.D.C., E.H.S., E.L.M. wrote the manuscript.

## Extended figures

**Extended Data Fig. 1:**
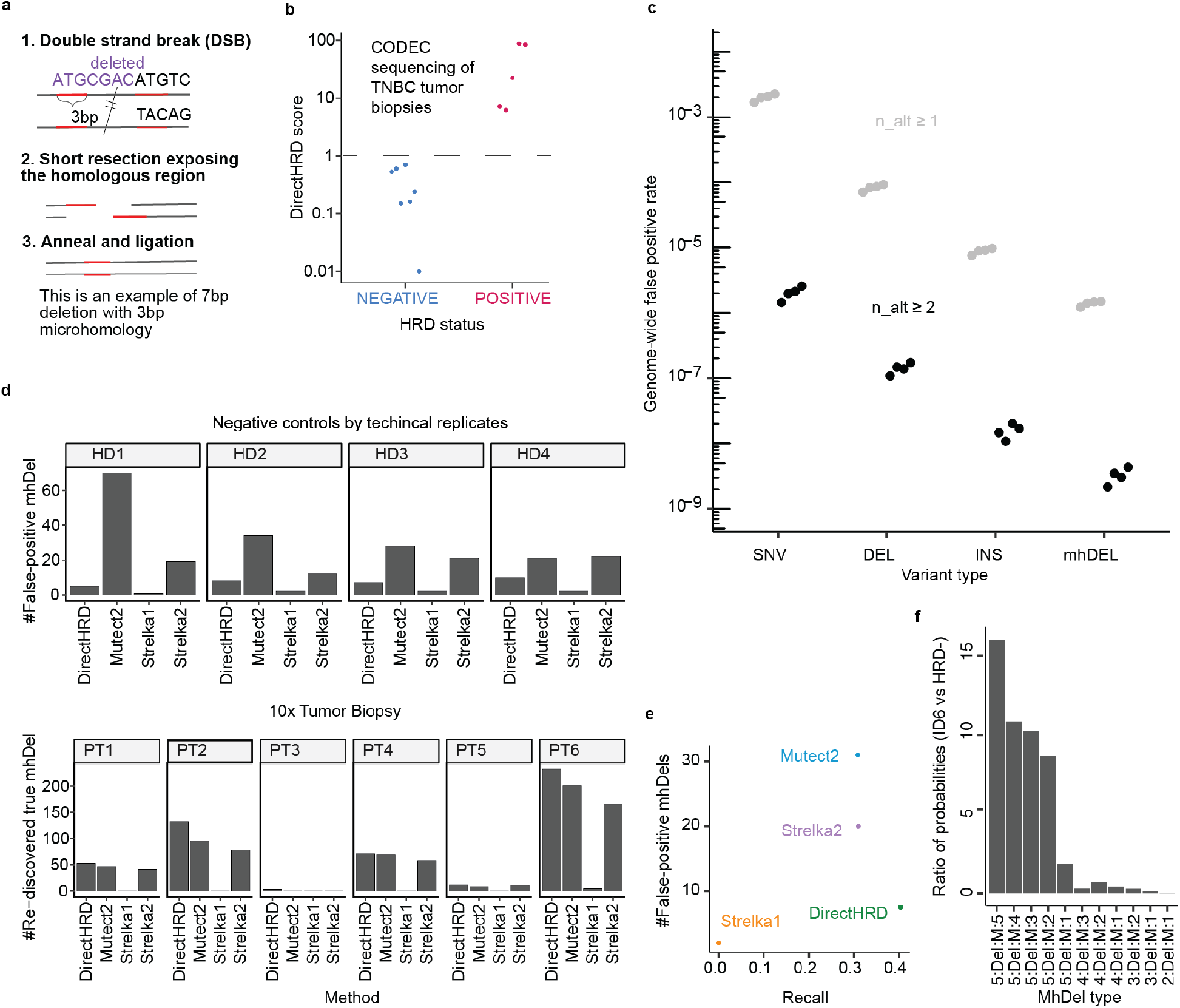
Optimized mhDel caller in DirectHRD. **a**, Example illustration of the creation of an mhDel by repairing a double strand break. **b**, DirectHRD scores of 12 TNBC tumor biopsies using CODEC^9^. Dashed line represented the HRD score cutoff 1, which correctly classified all samples based on Myriad MyChoice HRD testing as ground truth. **c**, Error rate of our Single Fragment Caller (SFC), used by DirectHRD, on 4 normal-normal pairs (negative controls, which were also used in **d,e**) from Illumina sequencing of different variant types. When requiring at least 2 unique fragments for calling a mutation, the error rate was several orders of magnitude lower than requiring at least 1 unique fragment. **d-e**, Performance of mhDel calling (DirectHRD, Mutect2, Strelka1 and Strelka2) on a set of 6 positive control samples (cancer patients, PTs) and 4 negative control samples (healthy donors, HDs). The recall is defined as the number of rediscovered mhDels in the 10x tumor WGS when the ground-truth set of mhDels were generated from 60x tumor WGS by taking the consensus results of Mutect2 and Strelka2. False-positive mhDels were called from two technical replicates of the same HD’s buffy coat DNA. **d**, Detailed numbers of true positives and false positives for each individual and each method. **e**, The median recall and the median number of false positive mhDels per method. **f**, The 11 mhDel types used in the DirectHRD model. Each mhDel type appears with a different probability in the HRD+ (ID6) signature vs HRD-signatures. The ratios between the two probabilities support HRD positivity when they exceed a value of 1. The label/definition of these mhDel types can be found in https://cancer.sanger.ac.uk/signatures/id/

**Extended Data Fig. 2:**
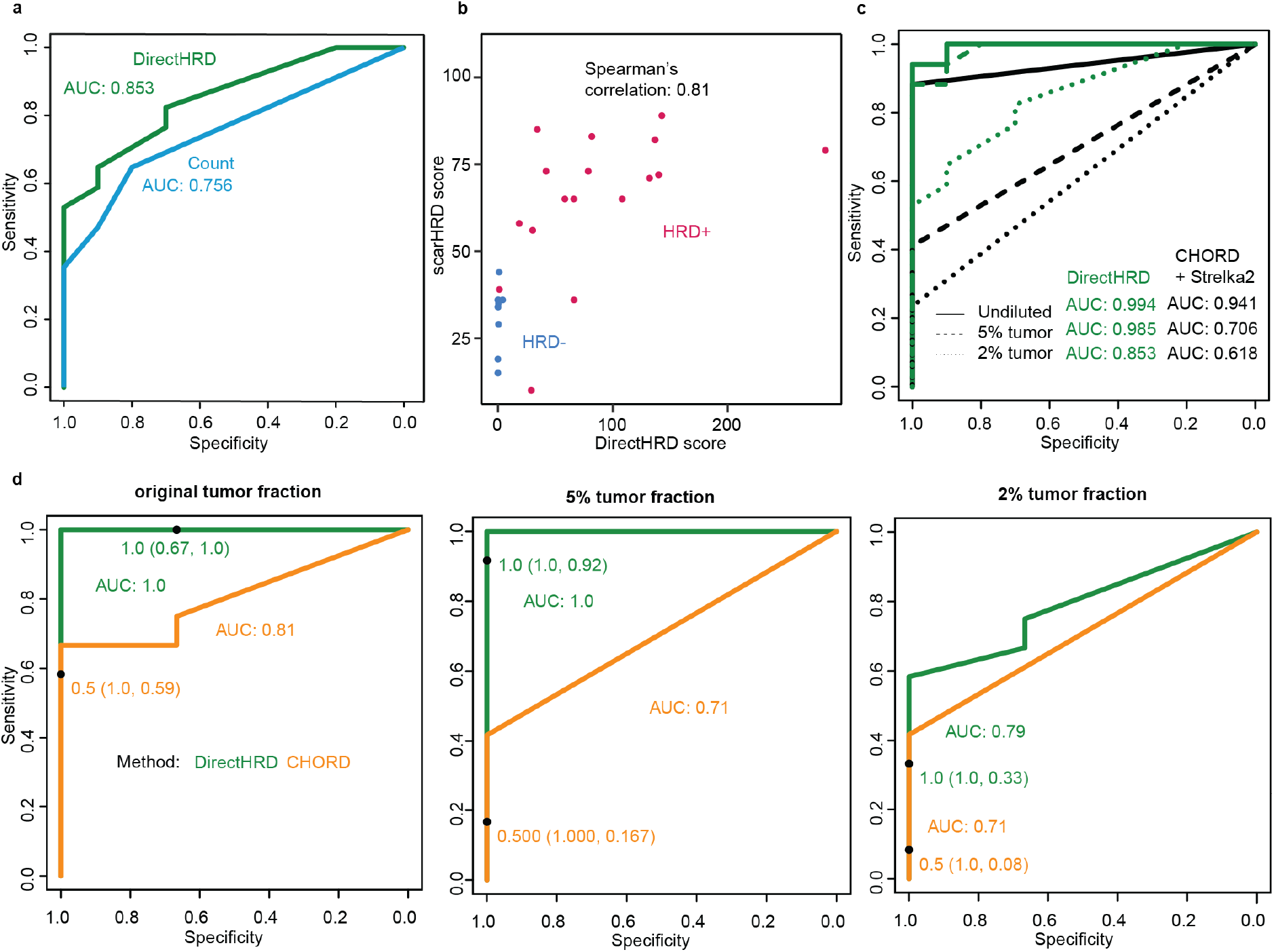
Additional DirectHRD results on tumor biopsies in stage II-III breast cancer (TNBC) cohort. **a**, Comparison of DirectHRD’s classification power using a probabilistic model to a count model at 2% tumor fraction for the in-silico-diluted tumor biopsy samples. The count model is simply using the number of mhDels with ≥2 microhomology (feature used in CHORD) for classification. **b**, Scatter plot of DirectHRD scores vs scarHRD scores using the 27 tumor biopsies at 15x, colored by HRD status from the Myriad MyChoice assay. Spearman’s correlation is shown. **c**, Comparison of DirectHRD vs CHORD *j* Strelka2 on 15x tumor biopsy samples and the in silico dilutions at 5% and 2% tumor fractions, also at 15x. **d**, Replot of performance of DirectHRD vs CHORD in Fig. 1e using only 15 TNBC tumor samples (3 were HRD-negative, 12 were HRD-positive) after removing the tumor samples also sequenced by CODEC (n=12). Receiver operating characteristic (ROC) curves, area under the curve (AUC), sensitivity and specificity (black dots) when using default cutoffs for HRD detection (DirectHRD HRDscore: 1.0, CHORD p-value: 0.5) are shown for original, 5% and 2% tumor fractions, all at 15x coverage.

**Extended Data Fig. 3:**
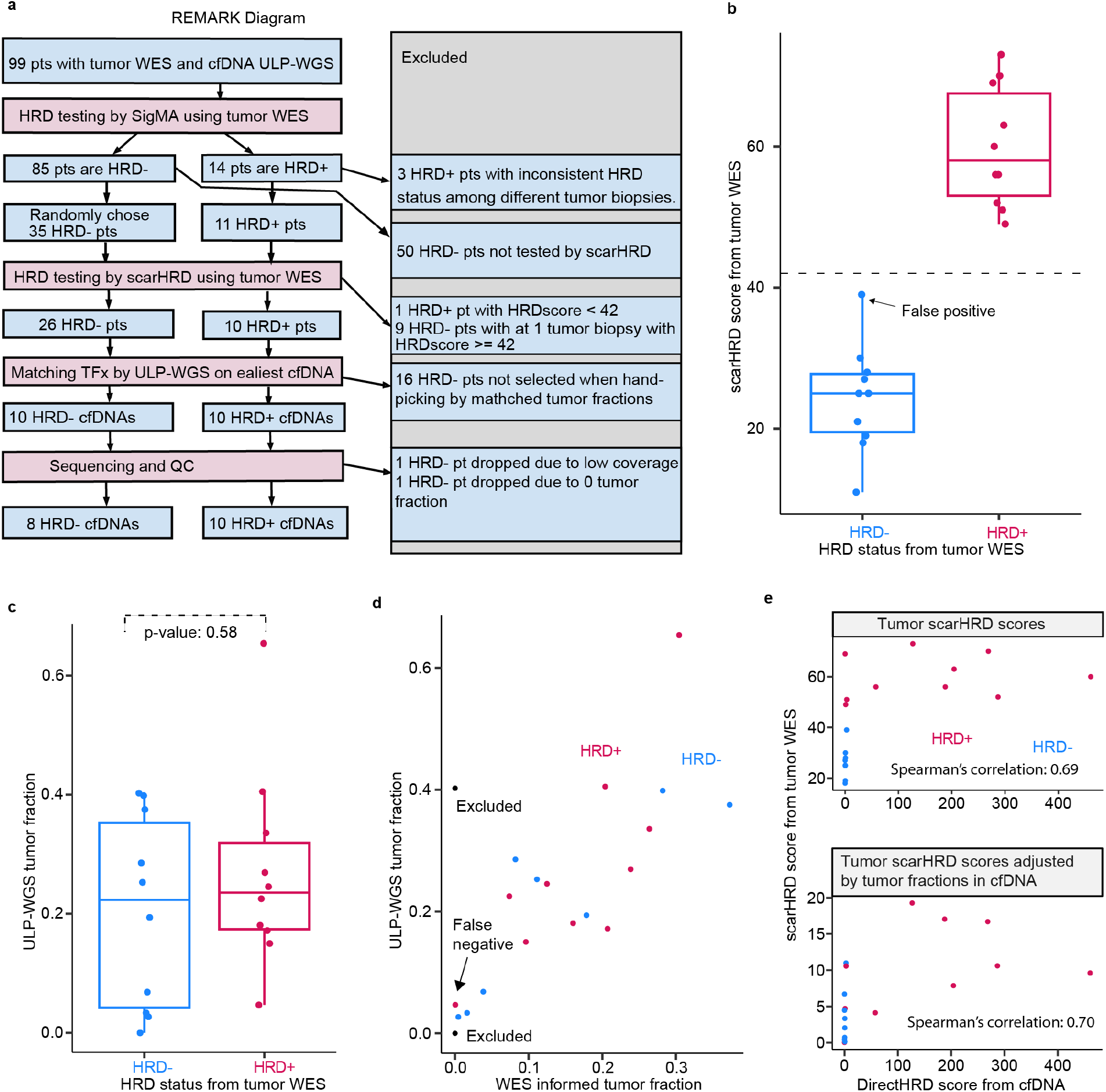
Stage IV (Metastatic) breast cancer cohort. **a**, REMARK diagram. pt: patient; TFx: tumor fractions. **b**, Scatter plot overlaid by box plot of 20 scarHRD scores, separated by HRD status inferred from WES of tumor biopsies by consensus results of scarHRD and SigMA. The cutoff value of 42 used here is indicated by a horizontal dashed line. **b,d**,The false positive and false negative cfDNA samples by DirectHRD were shown (ground*-*truth HRD status by tumor WES, see **a**). **c**, Scatter plot overlaid by box plot of tumor fractions by ichorCNA of 20 ULP-WGS libraries that were selected for deeper WGS, separated by HRD status inferred from tumor WES. Mann–Whitney U test p-value showed no statistical*ly significant* difference between the two distributions. **b,c**, Center lines, boxes, and whiskers indicate medians, 25% and 75% percentiles, and 5% and 95% percentiles, respectively. **d**, Tumor fractions estimated from previous ULP-WGS cfDNA samples using ichorCNA vs re-estimation from deeper WGS using tumor WES*-*informed SNVs. *Two* out of the 20 samples were excluded *from* further analysis because no tumor DNA was found in deeper WGS (pt300: ULP-WGS: 0.4, WES informed: 0; pt306: ULP-WGS: 0, WES-informed: 0), and one of the two samples also had low coverage (pt300: 4x). **e**, Scatter plot of DirectHRD scores from cfDNA vs scarHRD scores from tumor WES in 18 patients (2 excluded from **d**). The scores from the tumor samples are shown as both unadjusted (top) and adjusted by tumor fractions in cfDNA (bottom). Spearman’s correlations are presented. (**b-e**) Color coding corresponds to the HRD status of the patient being positive (magenta) vs. negative (blue) by tumor WES.

**Extended Data Fig. 4:**
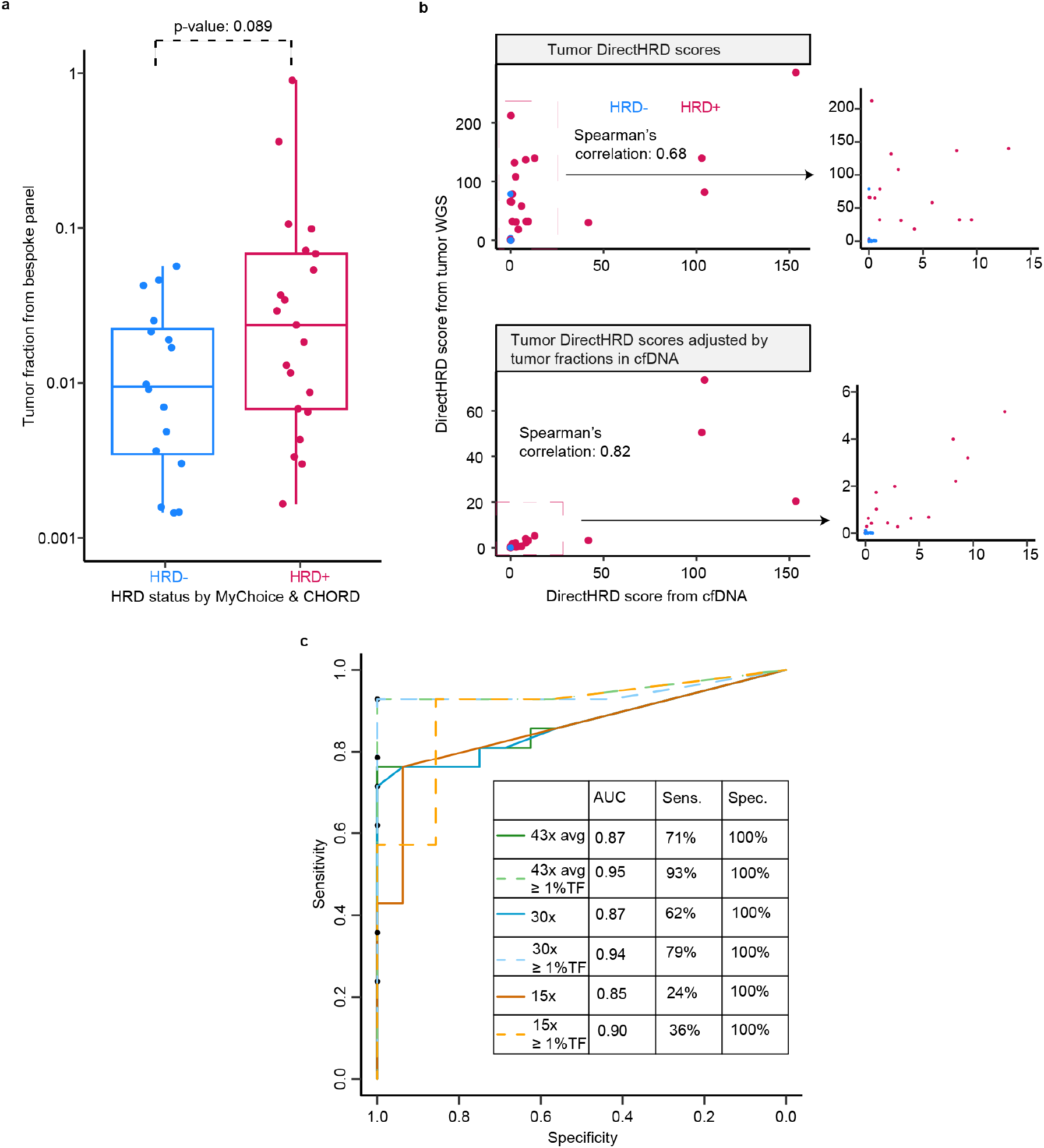
Additional DirectHRD results on cfDNAs in stage II-III breast cancer (TNBC) cohort. **a**, Scatter plot overlaid by box plot of estimated tumor fractions from personalized targeted panel sequencing of the same cfDNA samples by MAESTRO^15^, separated by HRD status from the Myriad MyChoice test (n=29) and CHORD (n=4) applied to the tumor biopsies. Center lines, boxes, and whiskers indicate medians, 25% and 75% percentiles, and 5% and 95% percentiles, respectively. Mann–Whitney U test p-value showed no statistical difference between the two distributions. **b**, Scatter plot of DirectHRD scores from cfDNA vs tumor WGS. The scores from tumor samples are shown as both unadjusted (top) and adjusted by tumor fraction in cfDNA (bottom). Highlighted regions are shown as zoom-ins on the right. Spearman’s correlations are presented. **a-b**, Color coding corresponds to the HRD status of the patient being positive (magenta) vs. negative (blue) from MyChoice testing of the tumor. **c**, Receiver operating characteristic (ROC) curves and tabulated results of area under the curve (AUC), sensitivity (Sens.) and specificity (Spec.) of DirectHRD on the 37 cfDNAs in the stage II-III breast cancer cohort at 3 different sequencing depths: median 43x (original, non-downsampled data), 30x and 15x. Additionally, we also evaluated the results when restricting only samples with ≥ 1% tumor fractions (dashed lines).

**Extended Data Fig. 5:**
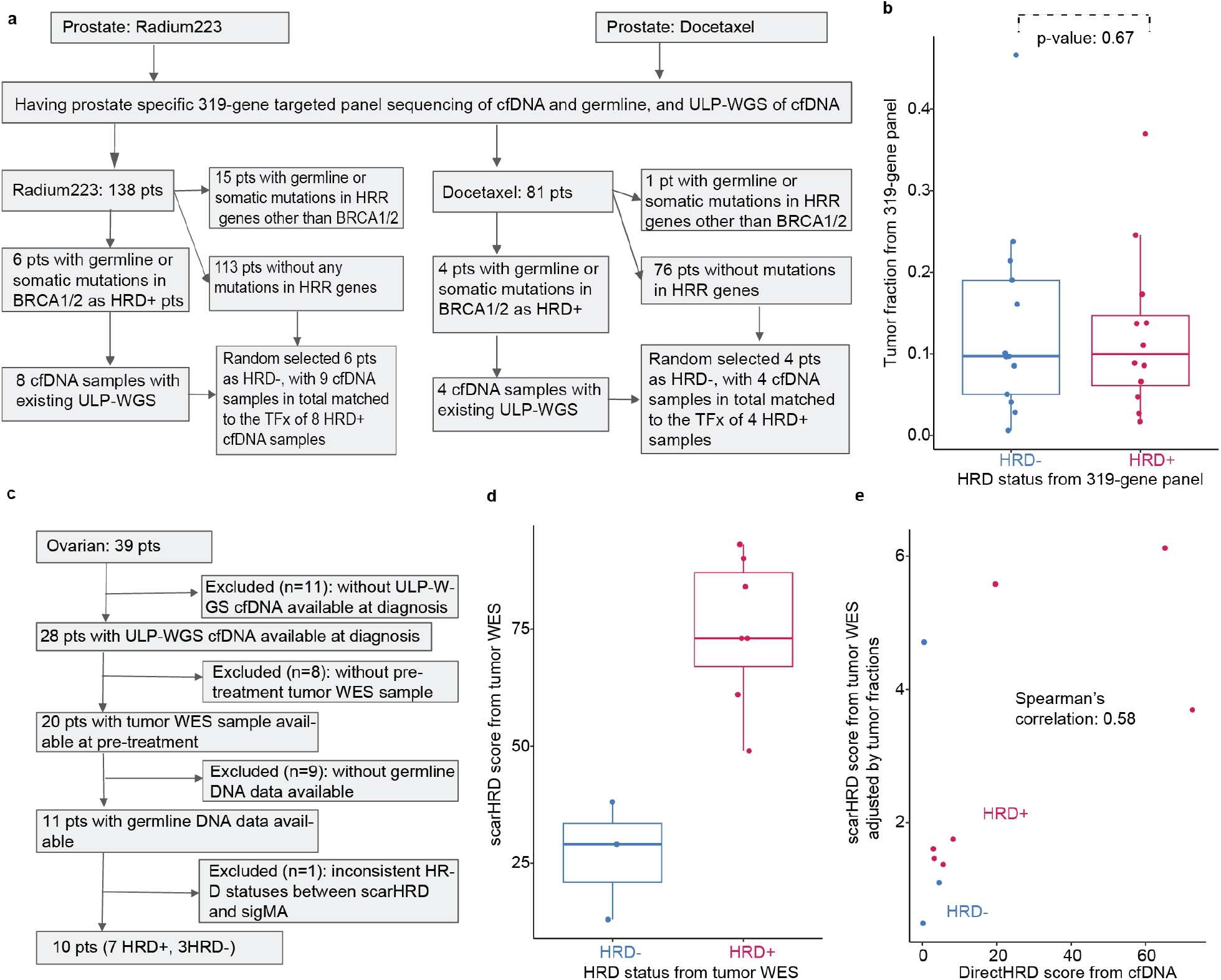
Two independent hold-out validation cohorts (Prostate: a, b; Ovarian: c, d, e): **a**, REMARK diagram of prostate cancer cohort (metastatic castration-resistant prostate cancer, mCRPC). pt: patient; TFx: tumor fractions; HRR: Homologous Recombination Repair. Patients who were HRD-positive had pathogenic or likely pathogenic germline or somatic mutations in BRCA1/2 genes (by ClinVar categories 4/5) by prostate cancer specific panel sequencing of patients’ cfDNA and germline samples. Patients who were HRD-negative had no pathogenic or likely pathogenic germline or somatic mutations in any of the HRR genes. **b**, Scatter plot overlaid by box plot of tumor fractions, separated by HRD status inferred from 319-gene targeted panel. Mann–Whitney U test p-value showed no statistical difference between the two distributions. **c**, REMARK diagram of ovarian cancer cohort. **d**, Scatter plot overlaid by box plot of scarHRD scores of the 10 ovarian cancer patients, separated by HRD status inferred from WES of tumor biopsies by consensus results of scarHRD and SigMA. **b,d** Center lines, boxes, and whiskers indicate medians, 25% and 75% percentiles, and 5% and 95% percentiles, respectively. **e**, Scatter plot of DirectHRD scores from cfDNA vs scarHRD scores from tumor WES in 10 ovarian patients. The scores from the tumor WES samples are shown as adjusted by tumor fractions in cfDNA. Spearman’s correlations are presented. (**b,d,e**) Color coding corresponds to the HRD status of the patient being positive (magenta) vs. negative (blue) by tumor WES.

