## Supplemental Table Info for "DirectHRD enables sensitive scar-based classification of homologous recombination deficiency (HRD)"

Supplementary Tables

1. Stage II-III breast cancer cohort patients and samples information
2. PCAWG analysis results
3. Stage IV breast cancer cohort patients and samples information
4. Stave IV prostate cancer cohort patients and samples information
5. Stave III-IV ovarian cancer cohort patients and samples information
6. DirectHRD results
7. CHORD results
8. Performance of DirectHRD on first available liquid biopsies from stage II-III breast cancer patients with MyChoice results.
9. Performance of DirectHRD on liquid biopsies from stage II-III breast cancer patients whose tumors had not been sequenced by CODEC
